# Pan-cancer analysis of whole genomes

**DOI:** 10.1101/162784

**Authors:** Peter J Campbell, Gaddy Getz, Joshua M Stuart, Jan O Korbel, Lincoln D Stein on behalf of the ICGC/TCGA Pan-Cancer Analysis of Whole Genomes Network

## Abstract

We report the integrative analysis of more than 2,600 whole cancer genomes and their matching normal tissues across 39 distinct tumour types. By studying whole genomes we have been able to catalogue non-coding cancer driver events, study patterns of structural variation, infer tumour evolution, probe the interactions among variants in the germline genome, the tumour genome and the transcriptome, and derive an understanding of how coding and non-coding variations together contribute to driving individual patient's tumours. This work represents the most comprehensive look at cancer whole genomes to date. NOTE TO READERS: This is an incomplete draft of the marker paper for the Pan-Cancer Analysis of Whole Genomes Project, and is intended to provide the background information for a series of in-depth papers that will be posted to BioRixv during the summer of 2017.

## Introduction

Cancer is the second most frequent cause of death worldwide, killing more than 8 million people every year and responsible for 1 in 7 deaths^1^. Globally, cancer deaths will increase by more than 50% over the coming decades, attributable to a number of driving forces: an ageing population in high income countries; increased exposure to carcinogens such as tobacco^2^, air pollution and asbestos^3^ in low- and middle-income countries; declining physical activity with concomitant rise in obesity worldwide; and the continued expansion of the human population. While prevention and treatment of competing causes of mortality, such as cardiovascular disease and infections, have led to major improvements in life expectancy, the gains for cancer mortality have been more modest. For many patients, surgery remains the only curative option, but once the tumour has spread from its original site, cure is often elusive. Nonetheless, in the last 20 years, our deepening understanding of the biology of cancer has enabled development of new therapeutics effective in a handful of cancers^4,5^ – it is this success that motivates the desire to systematically characterise cancer biology across all tumour types.

‘Cancer’ is a catch-all term used to denote a set of diseases characterised by autonomous expansion and spread of a somatic clone. To achieve this behavior, the cancer clone must modify multiple cellular pathways that enable it to disregard the normal constraints on cell growth, to modify the local microenvironment favoring its own proliferation, to invade through tissue barriers, to spread to other organs, and to evade immune surveillance^6^. No single cellular programme directs these behaviors. Rather there are many different potential abnormalities from which individual cancers draw their own combinations. In that sense, the commonalities of macroscopic features across tumours belie a vastly heterogeneous landscape of cellular abnormalities.

This heterogeneity arises from the fundamentally stochastic nature of Darwinian evolution; a process that operates in somatic cells as much as species. The preconditions for Darwinian evolution are three: characteristics must vary within a population; this variation must be heritable from parent to offspring; and there must be competition for survival within the population. In the context of somatic cells, heritable variation arises from mutations acquired stochastically throughout life, notwithstanding potential additional contributions from heritable epigenetic variation. A subset of these mutations drive alterations in cellular phenotype, and a small subset of those variants confer an advantage on the clone in its competition to escape the tight physiological controls wired into somatic cells. The mutations conferring selective advantage on the clone we call ‘driver mutations,’ as opposed to the selectively neutral, or possibly slightly deleterious, ‘passenger mutations.’

The discovery that cancers carry recurrent and specific genetic abnormalities in the 1970s^7^ and early 1980s^8,9^ has fuelled four decades of research to define the catalogue of genes and mutations that can drive cancer. This has been accelerated by technological advances in genomic analysis, from gross descriptions of chromosome structure by chromosomal banding^7^ and other cytogenetic techniques, through positional cloning of inherited cancer genes^10^, low-throughput capillary sequencing^11^ and comparative genomic hybridisation^12^, to the current era of massively parallel whole genome sequencing^13–17^. The ever more populous catalogue of cancer genes has opened new therapeutic opportunities, with effective drugs being developed for the *BCR-ABL* fusion gene of chronic myeloid leukaemia, *ERBB2* amplifications of breast cancer and the *BRAF* point mutations of melanoma^18–20^, amongst others.

## International collaborations to sequence whole cancer genomes

The advent of massively parallel sequencing promised a future in which the cancer genome was finite and knowable. Early studies showed it was in theory feasible to document every somatic point mutation in a given cancer, every copy number change and every structural variant^14,15^. In 2008, recognising the opportunity this advance in technology provided, the global cancer genomics community established The International Cancer Genome Consortium (ICGC) with the goal of systematically documenting the somatic mutations found in 25,000 samples representing all common tumour types^21^.

The ICGC comprises researchers from The Cancer Genome Atlas (TCGA) in the USA plus those from 17 countries and other jurisdictions in Europe, Asia and the Americas. Each ICGC project is organised around a single tumour type or a set of related types, for which a set of tumour/normal pairs derived from a target of 500 donors were characterised by whole genome sequencing, exome sequencing, transcriptome and/or DNA methylation analysis. The sample size was chosen to provide enough power to detect significantly mutated genes in at least 3% of patients based on an initial estimate of the background mutation rate.

ICGC samples have been carefully pre-screened by histopathologists and clinicians in order to ensure the accuracy of diagnosis and quality of the sample. Sequencing of both tumour and matched constitutional DNA are required to meet minimum coverage and quality requirements. Following the precepts established in the Human Genome Project, data from ICGC are rapidly released to the wider scientific community under appropriate safeguards to ensure ethical and regulatory compliance^22^. Since 2008, funding for ICGC projects has amounted to more than USD$900,000,000, with individual funding commitments in some countries being the largest biomedical grants they had ever awarded.

To date, there are 90 ICGC projects, of which 76 have submitted data across 21 primary organ sites and 31 distinct tumour types. At the time of writing, genomic data from 20,343 individual cancer patients were registered in the Data Coordination Center (https://dcc.icgc.org/), of whom 17,570 have molecular data, mostly exomes. Many major breakthroughs in the biology of individual tumour types have emerged from these studies, too numerous to cite exhaustively here, but including discoveries of new cancer genes and pathways^23–27^; insights into the underlying mutational processes operative in human cancers^28–32;^ delineation of the patterns of tumour heterogeneity and clonal evolution^33–36^; and development of genomics approaches to inform cancer prevention^37^ and clinical management of patients with cancer^38–40^. Many of these discoveries were enabled by novel computational and statistical methods designed to accurately detect various genomic alterations from sequencing data and analyse them across cohorts of patients to extract new biological insights.

## The Pan-Cancer Analysis of Whole Genomes Collaboration

The early studies from ICGC and TCGA revealed both commonalities and differences of somatic genomic architecture across tumour types. Some cancer genes are mutated in many different tumour types; others are specific to a single histological subtype^41,42^. All common tumour types are characterised by few frequently mutated genes and many rarely mutated genes; the patterns of co-mutation result in a huge diversity of combinations of driver mutations across individual patients^43–45^. Some tumours are driven by coding point mutations while others evolve through large-scale restructuring of chromosomes^46^; some cancer types mutate predominantly tumour suppressor genes^47^ while others have high frequency of driver mutations activating oncogenes^48^.

Numerous studies point to the relevance of non-coding regions, and projects including ENCODE,^49^ Blueprint^50^ and Epigenome Roadmap^51^ have revealed extensive catalogues of tissue-specific regulatory elements. Transcription factors and other proteins interact with enhancers, silencers, boundary elements, and overall chromatin structure to confer cell-specific regulatory responses, and recent studies have revealed the relevance of this interplay in cancer.^52-57^ Given that cells are pre-wired according to built-in control logics that involve coding and non-coding components, it stands to reason that changes in the DNA that affect these factors may underlie the tissue-specific nature of cancer onset and progression. Indeed, some evidence points this way, for example there is evidence that epigenetic marks are associated with mutation densities in cancer,^58,59^ whereas cancer-risk associated germline variants typically occur in intergenic regions and show enrichment with enhancers.^60^

The large number of samples subjected to whole genome sequencing by the ICGC now provides the opportunity to closely examine cancers beyond their protein-coding exomes, which are likely to provide only partial insights into the genomic landscape of cancer. Beyond providing insights into how mutations affect regulatory regions, whole genome sequencing can detail the full repertoire of classes of structural variation in cancers, facilitate resolving mutational processes and signatures acting in these, enable identifying viruses associated with cancers, and allow defining the full repertoire of germline variants in cancer patients. To tackle the various opportunties resulting from numerous cancer whole genome sequencing, 16 thematic Scientific Working Groups were formed and overseen by a Steering Committee for the PCAWG collaboration to pursue a multipronged analysis of the non-coding genome’s influence on cancer (Table 1).

**Table 1.** PCAWG Working Groups.

| Working Group Name | Role |
| --- | --- |
| PCAWG Technical Working Group | Clinical and molecular data management, execution of uniform alignment and core somatic mutation calling pipelines. |
| PCAWG Quality Control Working Group | Quality assurance |
| PCAWG Ethics and Legal Working Group | Policy and data access issues relating to distribution of data across compute clouds. |
| PCAWG Reference Annotations Working Group | Developed the reference set of genome annotations for use across the project. |
| PCAWG SNV Calling Working Group | Benchmarking of somatic single nucleotide variation and indel calling pipelines and development of methods for merging them. |
| PCAWG Drivers and Functional Interpretation Working Group | Identification and characterization of coding and non-coding drivers. |
| PCAWG Transcriptome Working Group | Exploration of the effect of genomic variation on transcription. |
| PCAWG Epigenome Working Group | Exploration of the interactions between genomic variation and the epigenome. |
| PCAWG Structural Variation Working Group | Identification and characterization of structural variations in the cancer genome. |
| PCAWG Mutational Signatures Working Group | Characterization of exposures and other mutational processes acting on the cancer genome. |
| PCAWG Germline Cancer Genome Working Group | Large scale haplotyping of PCAWG donors. Exploration of the interaction between germline and somatic mutations. |
| PCAWG Pathology and Clinical Correlates Working Group | Harmonization of tumour types and clinical descriptions. Exploration of clinical significance of somatic and germline variation. |
| PCAWG Evolution and Heterogeneity Working Group | Characterisation of evolutionary history of cancer genomes. |
| PCAWG Portals and Visualisation Working Group | Developed software systems for community access to PCAWG data set and interpretation. |
| PCAWG Mitochondrial Genome and Immunogenomics Working Group | Exploration of variation affecting the mitochondrial genome and the immune system.. |
| PCAWG Pathogens Working Group | Discovery of the presence of pathogen DNA in tumour samples and interpretation of significance. |

The maturing of datasets from individual ICGC and TCGA working groups presented the opportunity to formalise a meta-analysis of whole cancer genomes. However, algorithms for calling somatic mutations were not standardised among the different groups and had evolved considerably in the first few years of the consortium. For cross-tumour comparisons to be meaningful, the core bioinformatic analyses would need to be repeated using gold-standard, benchmarked, version-controlled algorithms. To achieve this, the Pan-Cancer Analysis of Whole Genomes (PCAWG) collaboration was established, comprising about 700 researchers from around the world. A Technical Working Group implemented the core informatics analyses, aggregating the raw sequencing data from the individual tumour type working groups, aligning it to the human genome and delivering a set of high quality somatic mutation calls for downstream analysis (**Figure 1**). Scientists from TCGA and ICGC submitted abstracts outlining potential research projects, which were aggregated into 16 thematic Scientific Working Groups. A Steering Committee oversaw the PCAWG collaboration, reporting to the executive committees of ICGC and TCGA.

**Figure 1.**
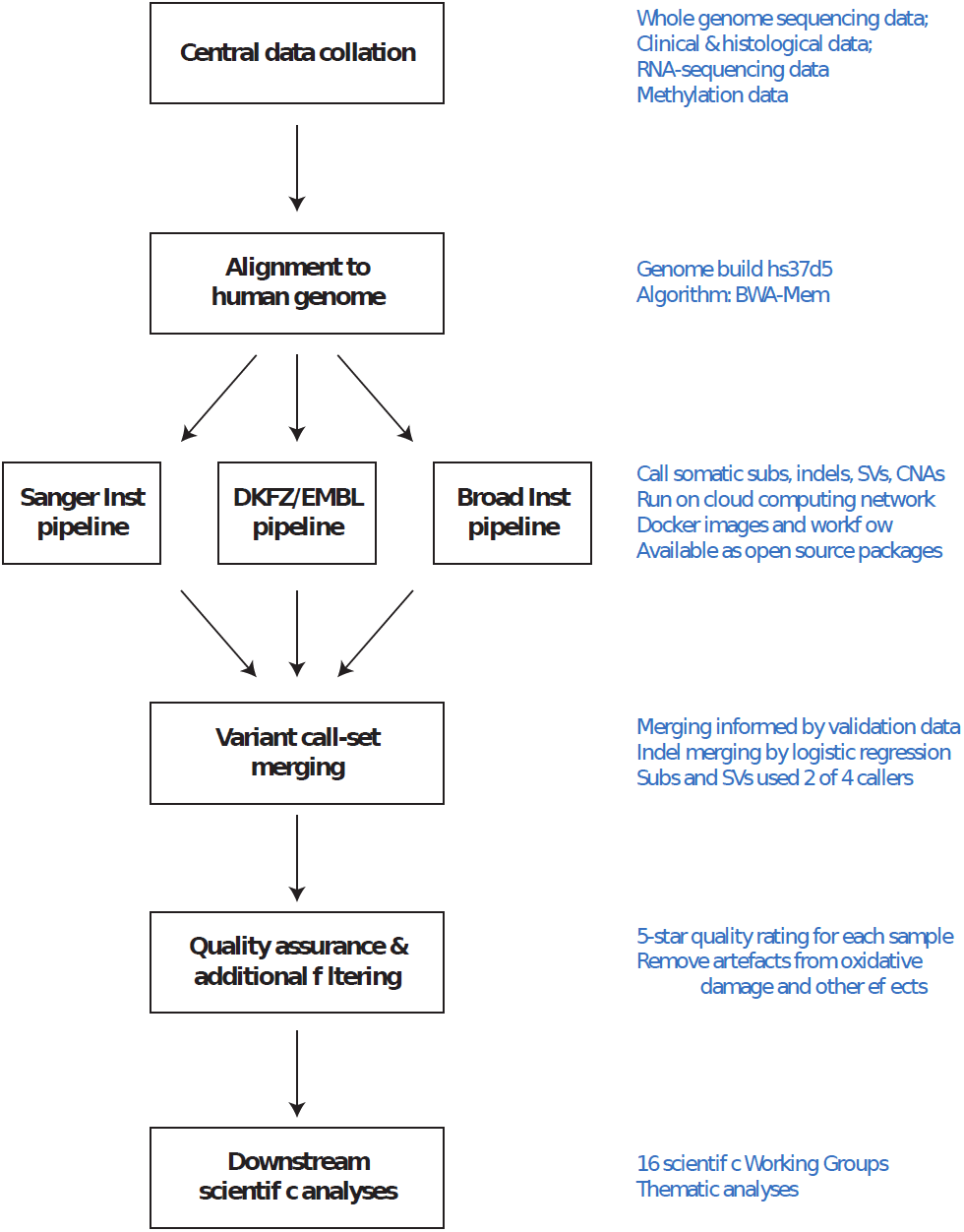
Flow-chart showing key steps in the analysis of PCAWG genomes

## Sample collection

Beginning in early 2015, we inventoried previous submissions of matched tumour/normal whole cancer genomes to the ICGC Data Coordinating Centre and polled ICGC projects for whole genomes that they anticipated completing in the near future. Our PCAWG inclusion criteria for donors included: a matched tumour and normal specimen pair; a minimal set of clinical information including patient age, sex and histopathological diagnosis; and characterisation of tumour and normal whole genomes using Illumina HiSeq platform 100-150bp paired-end sequencing reads. The minimum average depth required was 30 reads per genome base-pair in the tumour sample, and 25 in the normal sample. For the great majority of donors, the paired specimens consisted of a blood sample for the normal specimen, plus a fresh frozen sample of the primary tumour from a resection specimen. In a small number of cases the normal sample originated from tumour-adjacent normal tissue or another non-blood tissue (especially for blood cancers). Most of the tumour samples came from treatment-naïve, primary cancers, but there were a small number of donors with multiple samples of primary, metastatic and/or recurrent tumour. In addition to whole genome sequencing, roughly half of the donors had at least one tumour specimen that had been subjected to whole transcriptome analysis using RNA-sequencing, also centrally collected and re-analysed.

Ultimately, we collected genome data from a total set of 2,834 donors. After an extensive quality assurance process (described below), the data from 176 donors were deemed unusable and were excluded, leaving 2,658 donors, including 2,605 primary tumours and 173 metastases or local recurrences. Matching normal samples were obtained from blood (2,064 donors), tissue adjacent to the primary (87 donors), or other sites of normal tissue such as bone marrow, lymph node or skin (507 donors). The mean whole genome sequencing coverage in this set was 30 reads per base-pair for normal samples, while tumours had a bimodal coverage distribution with maxima at 38 and 60 reads per base-pair. For 75 donors, QA results were borderline and these donors were flagged in order to caution consortium members that they might be unsuitable for certain types of analysis, leaving a high-quality core of 2,583 donors. RNA-sequencing data was collected on 1222 donors with genome data, including 1178 primary tumours, 67 metastases or local recurrences, and 153 matched normal tissue adjacent to the primary tumour.

Demographically, the cohort included 1469 males (55%) and 1189 females (45%), with a mean age of 56 years (median 60 years; range 1-90 years). By using population ancestry-differentiated single nucleotide polymorphisms (SNPs) derived from the germline calls, we were able to estimate the population ancestry of each donor. The continental ancestry distribution was heavily weighted towards Europeans (77% of total) followed by East Asians (16%), as expected by large contributions from European, North American, and Australian projects (**Supplementary Table 1**).

## Histopathology harmonisation

In order to simplify the process of cross-tumour analyses, the PCAWG Pathology and Clinical Correlates Working Group consolidated and harmonised the histopathology descriptions of the tumour samples, using the icd-0-3 tumour site type controlled vocabulary (https://seer.cancer.gov/icd-o-3/) as its basis, in consultation with the leads of each of the contributing projects and a small group of expert anatomic pathologists. We described each tumour type using a four-tier hierarchical system consisting of Embryonic Origin (Mesoderm, Ectoderm or Endoderm), Organ System (such as Breast), Major Histologic Type (for example, Adenocarcinoma), and Major Histological Subtype (such as Infiltrating duct carcinoma). In addition, each tumour type was assigned a short abbreviation (*e.g.*, Breast-AdenoCa) and a standard colour for use in charts and tables. Overall, we established 39 distinct tumour types in the PCAWG data set (**Table 2**). The largest tumour type cohorts were hepatocellular carcinoma (Liver-HCC: 318 donors, 327 tumour specimens), pancreatic adenocarcinoma (Panc-AdenoCa: 239 donors, 241 specimens), and prostate cancer (Prost-AdenoCa: 210 donors, 286 specimens). Twelve tumour types had fewer than 20 representatives, including lobular carcinoma of the breast, cervical adenocarcinoma, and benign neoplasms of bone and cartilage. These tumour types, comprising a total of 56 specimens, were excluded from tumour-type specific cohort analyses due to lack of statistical power, but were included in pan-cancer analyses.

**Table 2.**
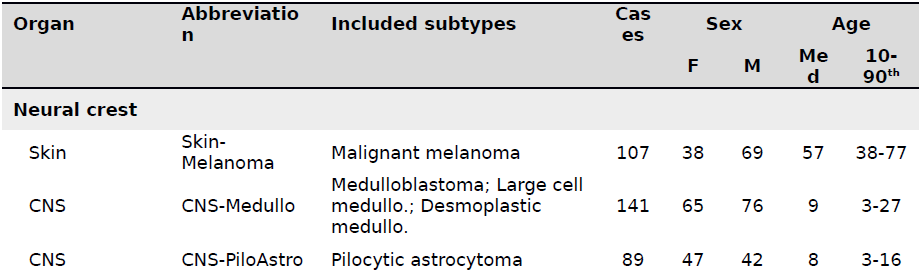
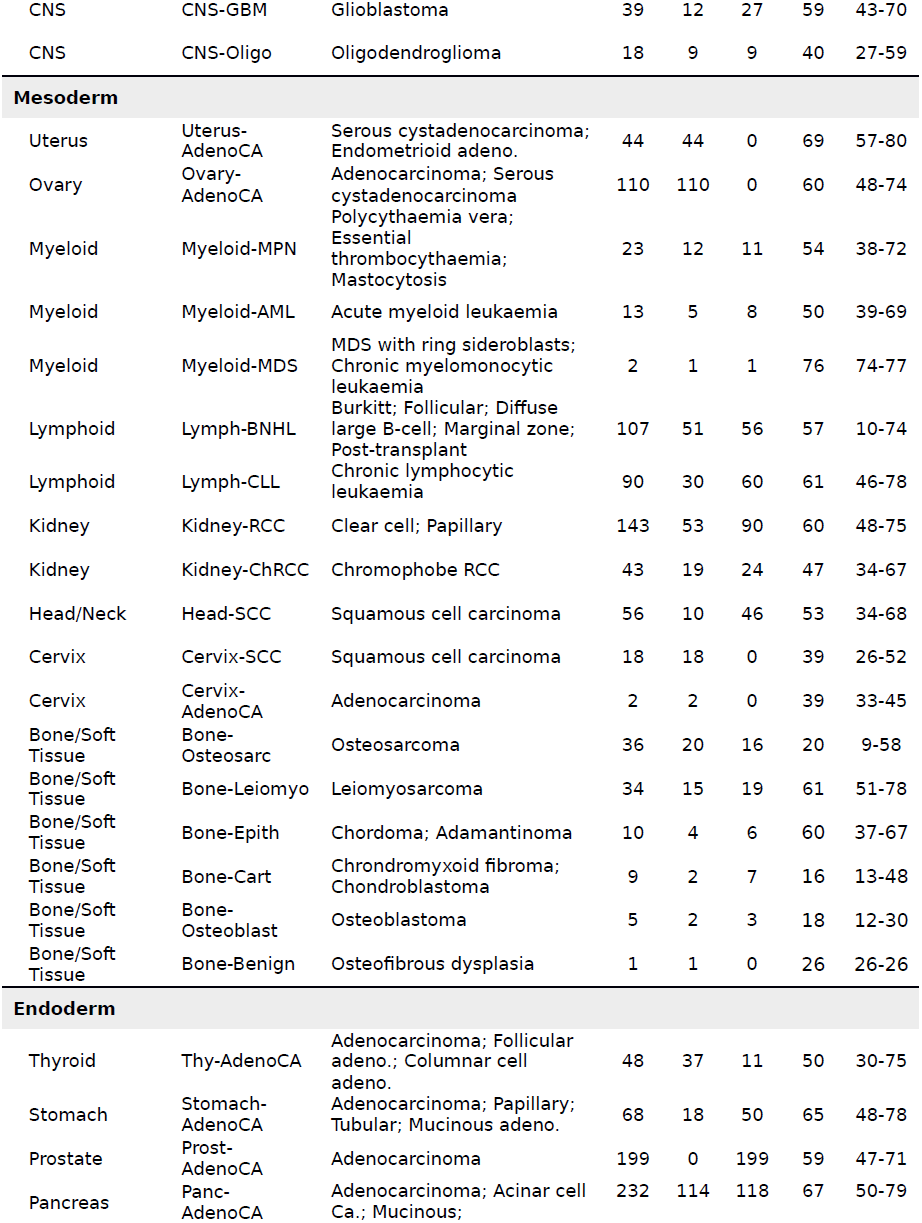
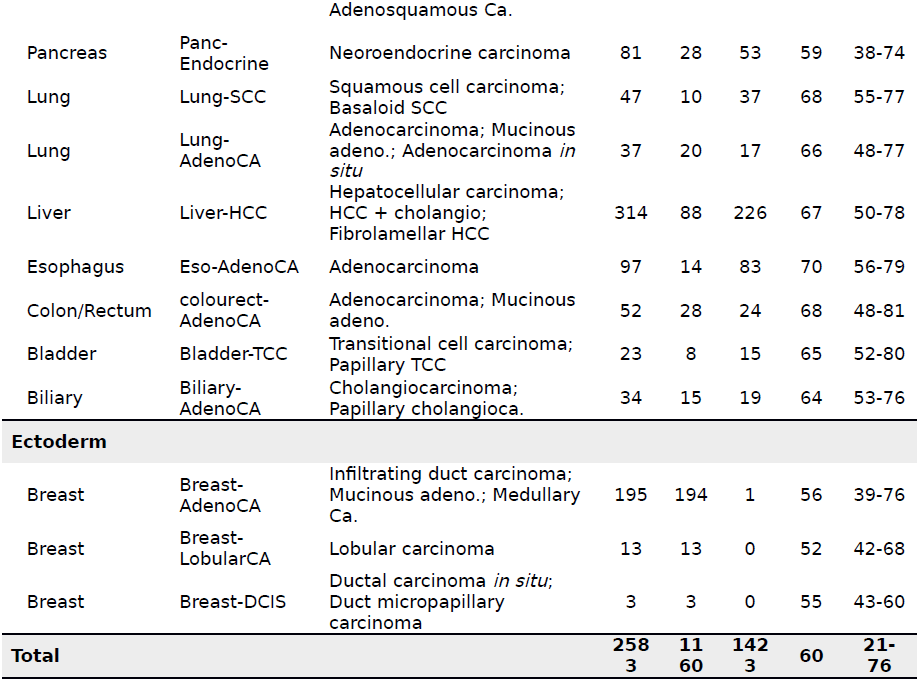
Overview of tumour types included in PCAWG project.

## Uniform processing and variant calling

In order to generate a consistent callset that could be used for cross-tumour type analysis, we analysed all samples using a uniform set of algorithms for alignment, variant calling, and quality control. We used the BWA-Mem algorithm^61^ to align each tumour and normal sample to human reference build hs37d50.^62^ Somatic mutations were identified in the aligned data using three established pipelines, run independently on each tumour/normal pair. Each of the three pipelines, labeled “Sanger”, “EMBL/DKFZ” and “Broad” after the computational biology groups that created and/or assembled them, consisted of multiple software packages for calling somatic single nucleotide variations (SNVs) and indels, copy number alterations (CNAs), and somatic structural variations (SVs). Each pipeline provided post-processing filters to remove likely false positive variant calls. A final set of filters were also run systematically across the entire set of PCAWG variants.

To assess the quality of the results from these three core pipelines, and to determine whether any other variant calling approaches would add additional value to the call set, we performed a systematic test and laboratory-based validation of 16 different computational pipelines. After this assessment, described below, we decided to run two additional callers^63,64^ on all samples to improve our ability to detect low-frequency SNVs and indels.

Following execution of each variant-calling pipeline, we merged the pipeline outputs for each variant type separately (SNVs, indels, CNAs, SVs) in order to achieve greater accuracy than provided by individual pipelines. The SNV and indel merge algorithms were designed and tested using the laboratory validation exercise described below as a gold standard.

RNA-Sequencing data were uniformly processed to produce normalised gene-level expression values, splice variant quantifications and measurements of alternative promoter usage, and to identify fusion transcripts, quantify allele-specific expression, and identify RNA edit sites. Calls of common and rare germline variants including single nucleotide variants, indels, SVs and mobile element insertions were generated using previously established principles for population-scale genetic polymorphism detection.^65,66^ The uniform germline data processing workflow comprised variant discovery using six different variant callers, followed by call-set merging, variant genotyping and statistical haplotype-block phasing. Somatic retrotransposition events, including *Alu* and LINE/L1 insertions,^67^ L1-mediated transductions^68^ and pseudogene formation,^69^ were called using a single, well-validated pipeline.^63^ We removed these retrotransposition events from the SV call-set. Mitochondrial DNA mutations were called using a published algorithm.^70^

## Core alignment and variant calling by cloud computing

The requirement to uniformly realign and call variants on more than 6,800 whole genomes presented significant computational challenges. The raw sequencing reads amounted to over 650 terabytes (TB), which corresponds to the size of a high definition movie running continuously for 30 years. If run serially, the execution of the alignment and the three variant-calling pipelines would have taken roughly 19 days/donor to execute on a single computer, or 145 years to complete the entire project. To accomplish this part of the analysis, we adopted a cloud-compute based architecture^71^ in which the alignment and variant calling was spread across 13 data centres distributed across three continents. The data centres represented a mixture of commercial infrastructure-as-a-service cloud compute, academic cloud compute, and traditional academic high-performance computer clusters, together contributing more than 10 million CPU core-hours to the effort. All told, the uniform alignment and variant calling took 23 months to execute – this included the data transfer, software development, and debugging time. On a cloud compute system running a fleet of 200 virtual machines, we estimate that without the overhead of software development and debugging, the project would take eight months to complete if repeated today.

The PCAWG-generated alignments, variant calls, annotations, and derived data sets are available for browsing and download at http://dcc.icgc.org/pcawg/. In addition, for the convenience of researchers who wish to avoid long data transfer times, a large subset of the data is pre-loaded and available for cloud-based computing on various platforms (see https://dcc.icgc.org/icgc-in-the-cloud).

## Quality assessment and control

Each donor and specimen was subject to a series of quality assessment (QA) and control (QC) steps. At the level of aligned reads, we tested for: minimum overall coverage of aligned reads; coverage across chromosomes; strand bias; insert size distribution; nucleotide content; base mismatch rate; indel rate; the number of unaligned reads; and concordance between the clinical sex of the donor and the sex inferred from the presence of Y chromosome markers and sex chromosome coverage. At the level of tumour/normal pairs and variant calls, we tested for: sharing of germline polymorphisms among the specimens from the same donor to detect sample swaps; the presence of common polymorphisms from two or more individuals to detect sample contamination; the presence of low-frequency somatic variants in the normal sample to detect tumour-in-normal contamination;^72^ and the presence of mutational signatures associated with sequencing artefacts such as oxidative damage. Of the 176 donors excluded on the basis of failing one or more of the QA tests, the most common reason for failure was RNA (cDNA library) contamination of tumour or normal, which manifested as multiple intron-length deletions in a substantial proportion of reads (39 donors). This was followed by lack of required clinical metadata, apparent misdiagnosis, or a disagreement between the clinical and genomic sex (29 donors), and unacceptably high levels of tumour DNA in the normal sample (15 donors). Sample swaps were relatively rare (6 donors), and there were a small number of donors excluded due to unique artefacts including contamination of tumour with a mouse library and the presence of a sibling’s genome in the blood of a leukaemia donor, presumably due to a bone marrow transplant. One cohort (of 33 acute myeloid leukaemias) was removed entirely due to a pervasive sequencing artefacts in SNV calls.

Among the non-excluded specimens, 735 showed signs of oxidative damage, as evidenced by high levels of G>T transversions among the variant calls.^73^ These artefactual variants were identified and removed by a purpose-built filter.^74^ The 75 donors that were deemed to be borderline following QA were flagged for a variety of reasons including an unexpectedly high fraction (>15%) of paired reads mapping to different chromosomes, an unusual mutational signature that did not correspond to a known biological process or artefact, or a level of tumour-in-normal contamination that approached, but did not exceed, the cut-off level (15%). We consider these suitable for some, but not all, analytic questions and left the choice of whether to use them or not to the downstream analytic groups.

## Validation, benchmarking and merging

In order to evaluate the performance of each of the mutation-calling pipelines and determine the strategy for integrating them, we performed a large-scale deep sequencing validation experiment. We selected a pilot set of 63 tumour/normal pairs from 23 cancer types across 26 contributing sequencing projects, on which we ran the three main mutation calling pipelines, and an additional 13 tools. The 63 tumours were chosen to have a wide range of somatic mutation frequencies in order to provide accurate representation of sensitivity and specificity estimates across samples. Of the 63 cases, 50 had sufficient DNA in both tumour and normal samples to enable deep sequencing targeting the putative mutated sites through DNA hybridisation capture. We selected ~250,000 SNVs and indels for validation by first stratifying mutations based on the number of methods that called them and then evenly sampling from each of these strata. This enabled us to estimate, for each method, false-positive and false-negative rates, which were used to calculate performance metrics such as precision, sensitivity and a combined (F1) score.

Next, we examined multiple methods for integrating calls made by each of the three pipelines. We evaluated the performance of simple methods (such as taking the union or intersection of the calls) as well as more sophisticated methods that used, beyond the three pipelines, additional parameters (such as coverage, variant allele frequency and nearby sequence context) to predict whether a mutation is real or not. The final consensus calls for SNVs were based on a simple approach that required two or more methods to agree on a call. For indels, because methods were less concordant, we used logistic regression to integrate the calls. The SV merge accepted all calls made by two or more of the four primary SV callers (one pipeline has two SV callers).

Overall, the sensitivity and precision of the consensus calls were 95% (CI_90%_: 88-98%) and 95% (71-99%) respectively for SNVs. For indels, in keeping with greater challenges in identification accuracy, sensitivity and precision were 60% (34-72%) and 91% (73-96%). Using manual assessment of called SVs as a gold standard, the false discovery rate of merged calls was estimated to be 2.5%, with 10% of true calls rejected. For all mutation types, accuracy was reduced in repeat-rich regions relative to coding and other unique regions.

## Pan-cancer burden of somatic mutations

Across the 2,583 donors in the PCAWG dataset, we called 43,778,859 SNVs; 410,123 somatic multi-nucleotide variants; 2,418,247 somatic indels; 288,416 SVs; 21,076 somatic retrotransposition events; and 8,185 *de novo* mitochondrial DNA variants (**Supplementary Table 1**). There was considerable heterogeneity in the burden of somatic mutations across patients and tumour types (**Figure 2**). For example, the median number of base substitutions across different tumour types spanned more than two orders of magnitude, from a median of 169/patient in pilocytic astrocytoma to 70,873/patient in melanoma. Similarly, within each tumour type, the burden of somatic substitutions typically varied over 2 orders of magnitude, with the range observed in breast adenocarcinoma being 1,203 in one patient to 65,065 in another. Similar heterogeneity was observed for other classes of somatic variation.

**Figure 2.**
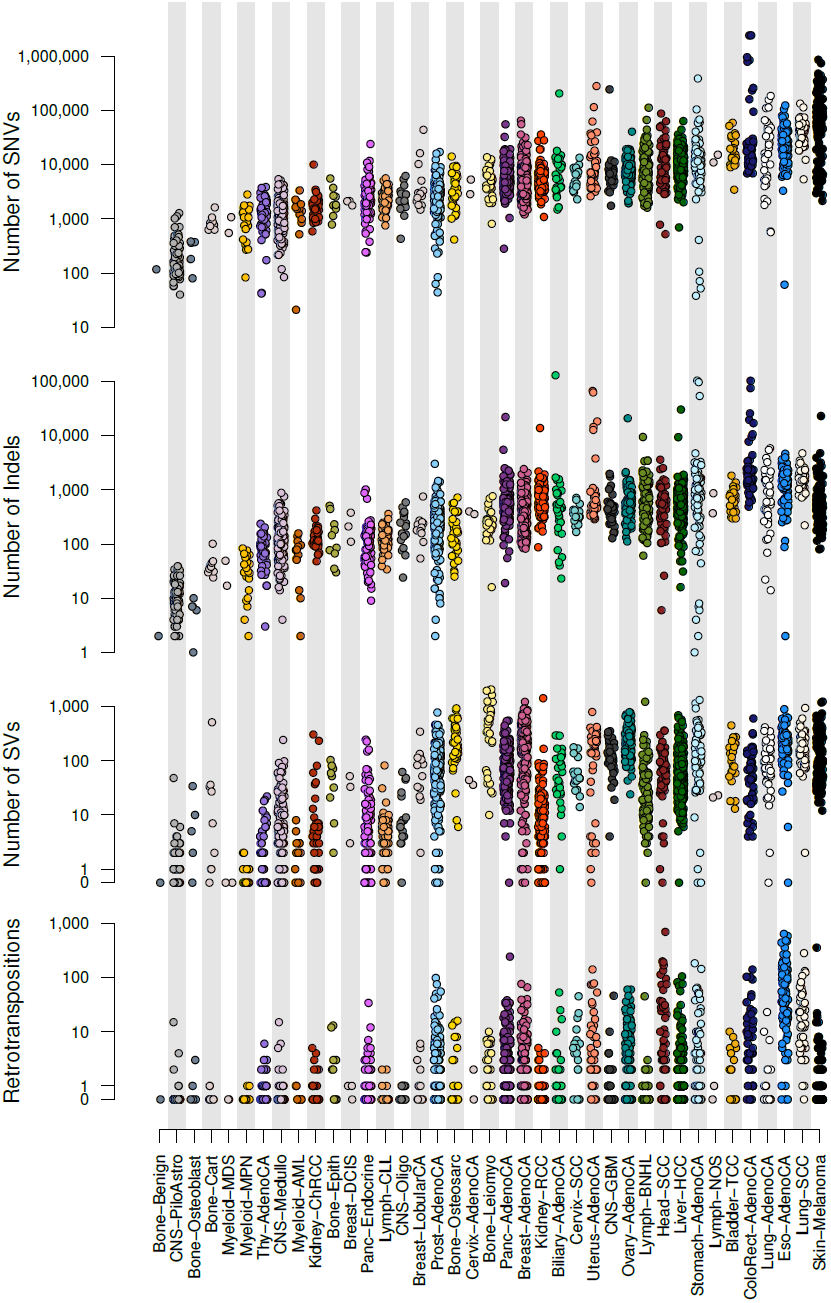
Distribution of numbers of somatic mutations of different classes across the different tumour types included in the PCAWG project. The y axis is on a log scale. SNVs, single nucleotide variants (single base substitutions); Indels, insertions or deletions <100 base pairs in size; SVs, structural variants; Retrotranspositions, counts of somatic retrotransposon insertions, transductions and somatic pseudogene insertions.

Strikingly, at the level of tumour types, there was a broad correlation in mutation burden among the different classes of somatic variation (**Figure 2**). Thus, melanomas, squamous cell carcinomas of the lung and oesophageal adenocarcinomas all showed high rates of somatic substitutions, indels, structural variation and retrotransposition. In contrast, the genomes of blood cancers and childhood brain tumours were generally quiet and stable, with relatively few variants of any type. Analysed at a per-patient level, this correlation held (**Supplementary Figure 1**).

This correlation in burden among different classes of somatic mutation has not been delineated on a pan-cancer basis before, and the underlying causes are unclear. It is likely that age plays some role – we observe acorrelation of most classes of somatic mutation with age at diagnosis (~190 substitutions/year, p=0.02; ~22 indels/year, p=5×10^-5^; 1.5 SVs/year, p<2×10^-16^; **Figure 3**). Other factors are also likely to contribute to the correlations among classes of somatic mutation, since there is evidence that some DNA repair defects can cause multiple types of somatic mutation^75^ and a single carcinogen can cause a range of DNA lesions.^76^

**Figure 3.**
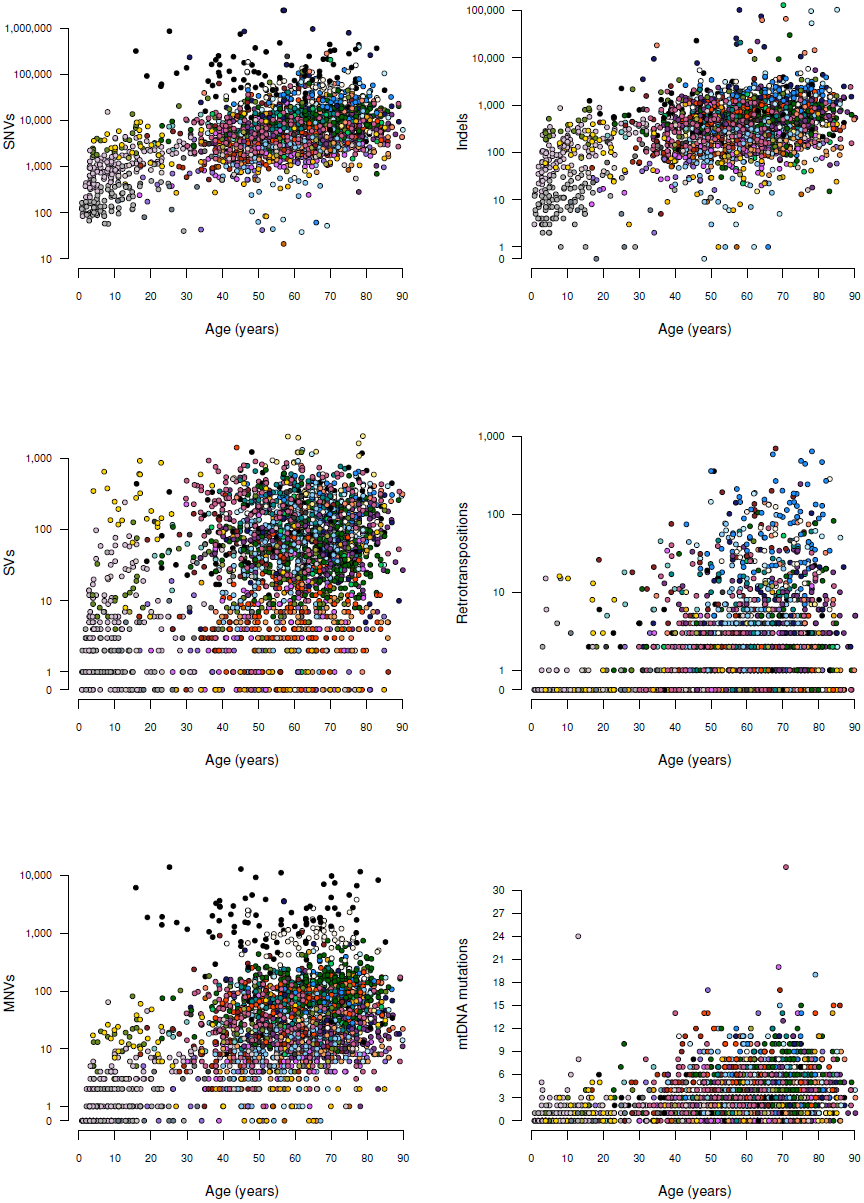
Numbers of somatic mutations by age at diagnosis. Points are coloured by tumour type, using the colour scheme in Figure 3. The y axis is on a log scale for all except mitochondrial DNA mutations. SNVs, single nucleotide variants (single base substitutions); Indels, insertions or deletions <100 base pairs in size; SVs, structural variants; Retrotranspositions, counts of somatic retrotransposon insertions, transductions and somatic pseudogene insertions; MNVs, multinucleotide variants (mostly dinucleotide substitutions); mtDNA mutations, number of somatic mutations in the mitochondrial genome.

## SUPPLEMENTARY FIGURE LEGENDS

**Supplementary Figure 1.**
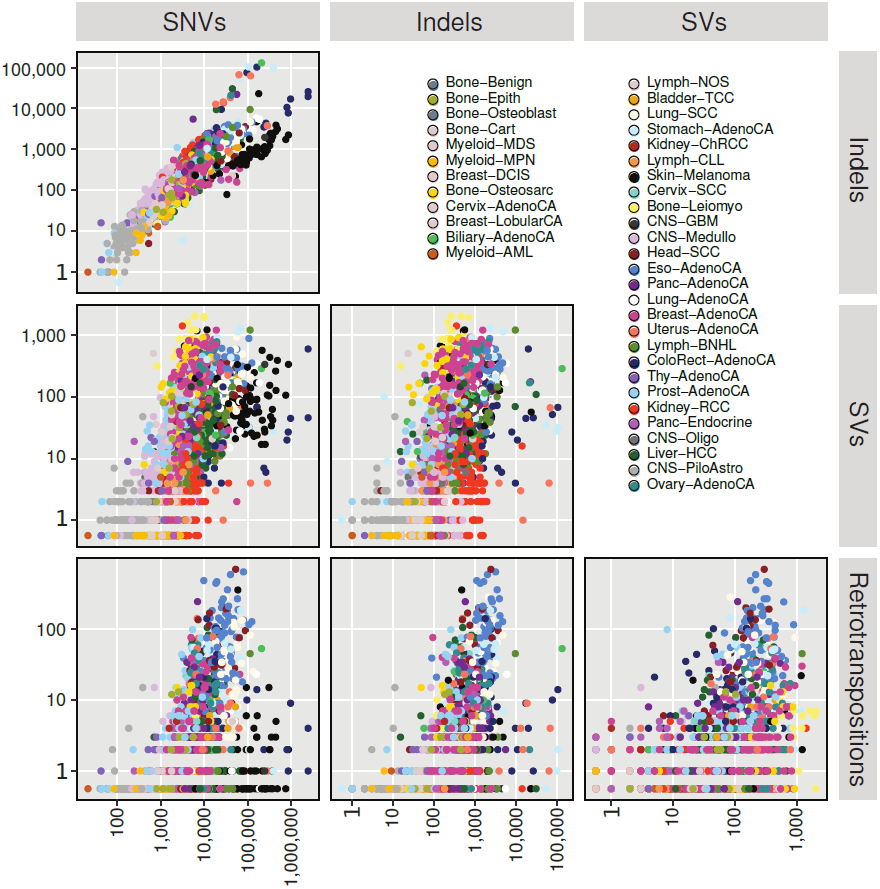
Pairwise comparison of rates of different classes of somatic mutation. Points are coloured by tumour type, as depicted in the legend. The y axes are on a log scale. SNVs, single nucleotide variants (single base substitutions); Indels, insertions or deletions <100 base pairs in size; SVs, structural variants; Retrotranspositions, counts of somatic retrotransposon insertions, transductions and somatic pseudogene insertions.

